# Curved flexible neural interfaces for laminar electrophysiology in deep-brain structures

**DOI:** 10.1101/2025.09.29.679383

**Authors:** Jo’Elen Hagler, Lucia Pizzoccaro, Yasser Matos-Peralta, Meijing Wang, Guillaume Ducharme, Bénédicte Amilhon, Fabio Cicoira

**Affiliations:** Department of Chemical Engineering, Polytechnique Montreal, Montreal H3T 1J4, Canada; Department of Neurosciences, University of Montreal, Montreal H3T 1J4, Canada; CHU Sainte-Justine Azrieli Research Center, Montreal H3T 1C5, Canada; Department of Engineering Physics, Polytechnique Montreal, Montreal H3T 1J4, Canada

## Abstract

Neural circuits are organized within complex three-dimensional architectures. Most neural interfaces follow linear insertion trajectories, limiting their ability to achieve laminar recording in brain regions where neuronal layers lie approximately parallel to the insertion path. Here, we introduce a curved flexible neural interface that enables near-perpendicular alignment of the recording sites with the targeted neuronal layers. The device consists of a flexible Parylene-C neural probe, integrating 16 PEDOT:BF_4_-coated Au microelectrodes and a transient silk fibroin stiffener for controlled implantation. The electrodes exhibit low impedance at 1 kHz (29.2 ± 2.5 kΩ) and were accurately positioned across the layers of the ventral hippocampus using a rotational implantation strategy. Chronic in vivo recordings demonstrate stable electrochemical performance and reliable acquisition of local field potentials over four weeks. This work establishes a strategy for anatomically matched neural interfacing, enabling high-resolution investigation of neural circuits that are challenging to study with conventional linear probes.

---

The intricate three-dimensional structure of the brain is fundamentally mismatched with the rigid, linear configuration of conventional deep-brain interfaces.^1^ Brain architecture is highly diverse, with structures such as the cortex, cerebellum and hippocampus, exhibiting a layered yet highly convoluted organization, while others display less ordered arrangements. Achieving a controlled orientation of linear micro-electrodes relative to the organization of neural tissue can be technically challenging and, in some cases, impossible. Overcoming this limitation is essential for applications requiring dense, laminar sampling of neuronal activity and local field potentials across neuronal layers that lie approximately parallel to conventional insertion trajectories.^2,3^ One such example is the hippocampus, a layered structure which follows a curved trajectory along the dorso-ventral axis of the brain. In the dorsal hippocampus, linear microelectrodes can readily be inserted orthogonally to the neuronal layers, an orientation required for laminar electrophysiological recording. Achieving an equivalent orientation in the ventral hippocampus is considerably more challenging, because conventional linear probes cannot readily follow its curved anatomy.^4,5^

Recent research in neural interfaces has largely focused on improving the structural, mechanical, and electrical properties of implantable devices.^6^ Flexible polymeric probes reduce the mechanical mismatch between neural tissue and implanted devices, thereby mitigating the chronic tissue response associated with rigid neural interfaces.^7–11^ However, despite these advances, flexible probes have largely retained conventional linear geometries. Likewise, conductive polymer coatings such as PEDOT have substantially reduced electrode impedance, enabling high-fidelity recording.^12–16^ Despite these advances, current neural interfaces remain predominantly based on linear architectures, leaving the challenge of achieving precise alignment with curved neuronal layers largely unresolved.

This study presents a strategy to overcome these geometric constraints by combining a flexible Parylene-C neural probe with a predefined curvature, a transient silk fibroin stiffener that provides temporary mechanical support during implantation, and a custom rotational manipulator to guide the device along a circular trajectory. The system targets the ventral hippocampus, a deep brain region whose layered anatomy is difficult to access using conventional laminar recording approaches.^17^ PEDOT:BF_4_-coated Au microelectrodes provide a low-impedance interface for chronic neural recordings while remaining compatible with the silk-assisted implantation strategy. Longitudinal in vivo recordings over four weeks demonstrate stable electrochemical performance and reliable acquisition of local field potentials across the ventral hippocampus. This work demonstrates that matching neural interface geometry to brain anatomy provides a practical strategy for laminar recordings in brain regions that are difficult to access using conventional linear probes

## Results

### System Design and Surgical Approach

The overarching goal of this work was to develop a neural interface capable of achieving orthogonal laminar recordings in deep brain structures with curved neuronal layers. To achieve this, we engineered an integrated system comprising a curved microelectrode array, a transient silk fibroin stiffener, and a custom rotational implantation device (Fig. 1).

**Fig. 1.**
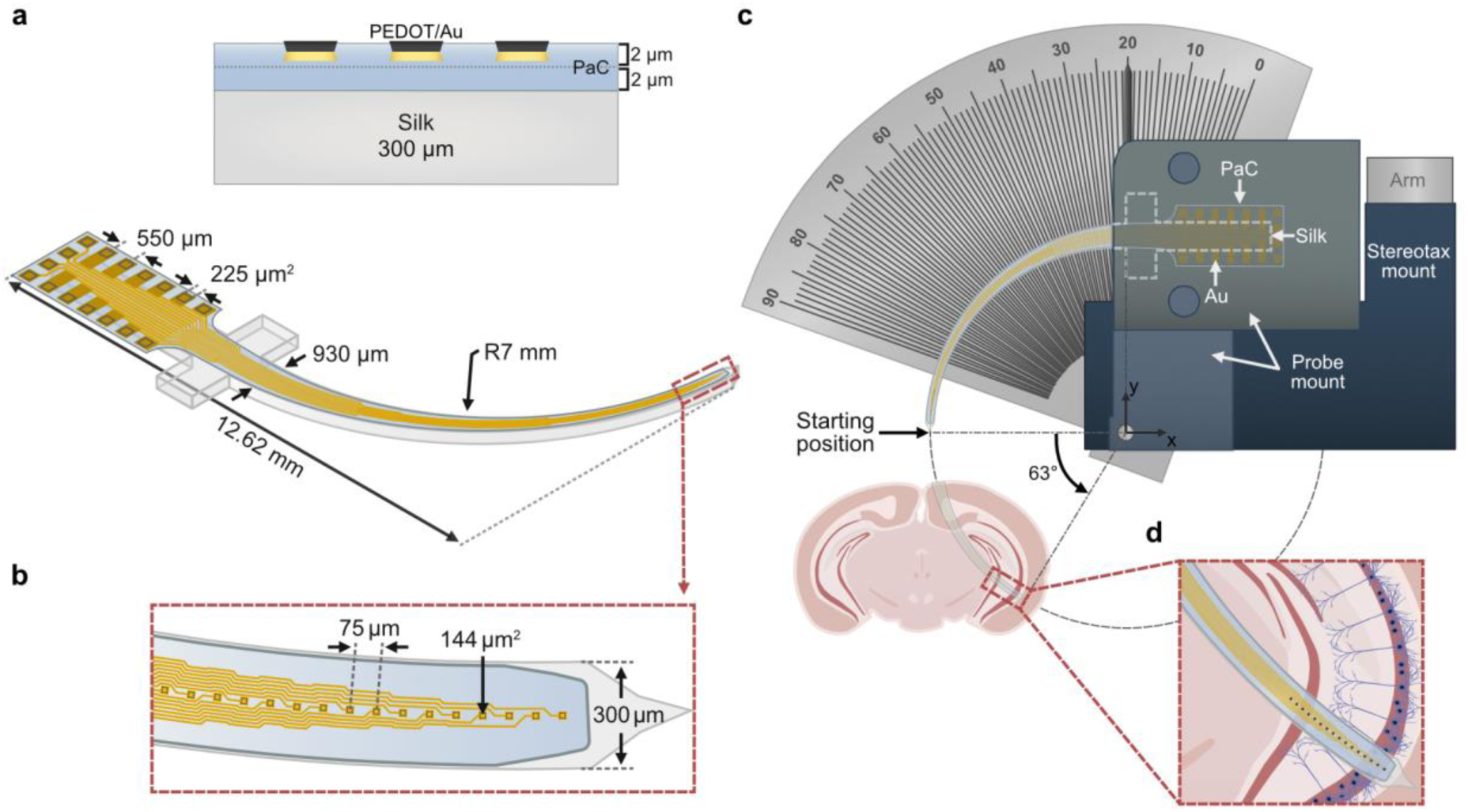
| Design, material architecture, and rotational implantation strategy of the curved microelectrode array. **a** 3D schematic and cross-sectional view of the multilayer neural probe, comprising PEDOT:BF_4_-coated Au microelectrodes encapsulated with a 4 μm-thick Parylene-C layer and supported by a 300 μm silk fibroin stiffener. **b** Magnified top view of the 16-channel array showing the sharpened probe tip and microelectrode layout optimized for high-density neural interfacing. **c** Schematic of the rotational implantation strategy. The probe is mounted on a custom rotational manipulator integrated with a stereotaxic frame, enabling insertion along a fixed 7 mm radius trajectory. The insertion angle is determined using a mouse brain atlas to target deep anatomical structures, such as the ventral hippocampus. **d** Schematic illustrating alignment of the curved array within hippocampal neuronal layers. The predefined curvature enables positioning parallel to the axodendritic axis of pyramidal neurons, enabling high-resolution laminar local field potential (LFP) recordings and current source density (CSD) analysis.

The neural interface consists of a flexible Parylene-C probe supported by a 300 μm-thick silk fibroin stiffener that provides the temporary mechanical rigidity required for implantation (Fig. 1a). The probe shank was designed with a 7 mm radius of curvature to match the anatomical trajectory of the ventral hippocampus (Fig. 1b). Each 16-channel array featured Au microelectrodes (≈144 μm^2^) coated with PEDOT:BF_4_ to reduce interface impedance and support high-fidelity signal acquisition.^12,18–20^ The stiffener is required because, although the relatively low Young’s modulus of Parylene-C (≈2.5 GPa) improves mechanical compliance with neural tissue,^21,22^ its low bending stiffness prevents direct insertion into deep brain regions.

The successful delivery of the curved probe was enabled by a custom-built rotational manipulator (Fig. 1c), which constrains the probe to a fixed circular trajectory. This strategy allows the recording sites to align orthogonally with target neuronal layers (Fig. 1d), an orientation required for laminar local field potential and high-resolution current source density (CSD) analysis.^23,24^

The fabricated device was first characterized by optical and scanning electron microscopy to verify the structural integrity of the integrated probe prior to electrochemical evaluation. Optical microscopy and scanning electron microscopy (SEM) confirmed the successful integration of the curved microelectrode array with the silk fibroin stiffener while preserving access to the recording sites (Fig. 2a). SEM images further revealed well-defined Parylene-C insulation around individual microelectrodes (Fig. 2b). Prior to PEDOT:BF₄ deposition, the Au electrode surface exhibited the characteristic granular morphology of evaporated thin films (Fig. 2c).^25^ Following electrodeposition, the PEDOT:BF_4_ coating displayed the characteristic nodular morphology (Fig. 2d).^26–28^

**Fig. 2.**
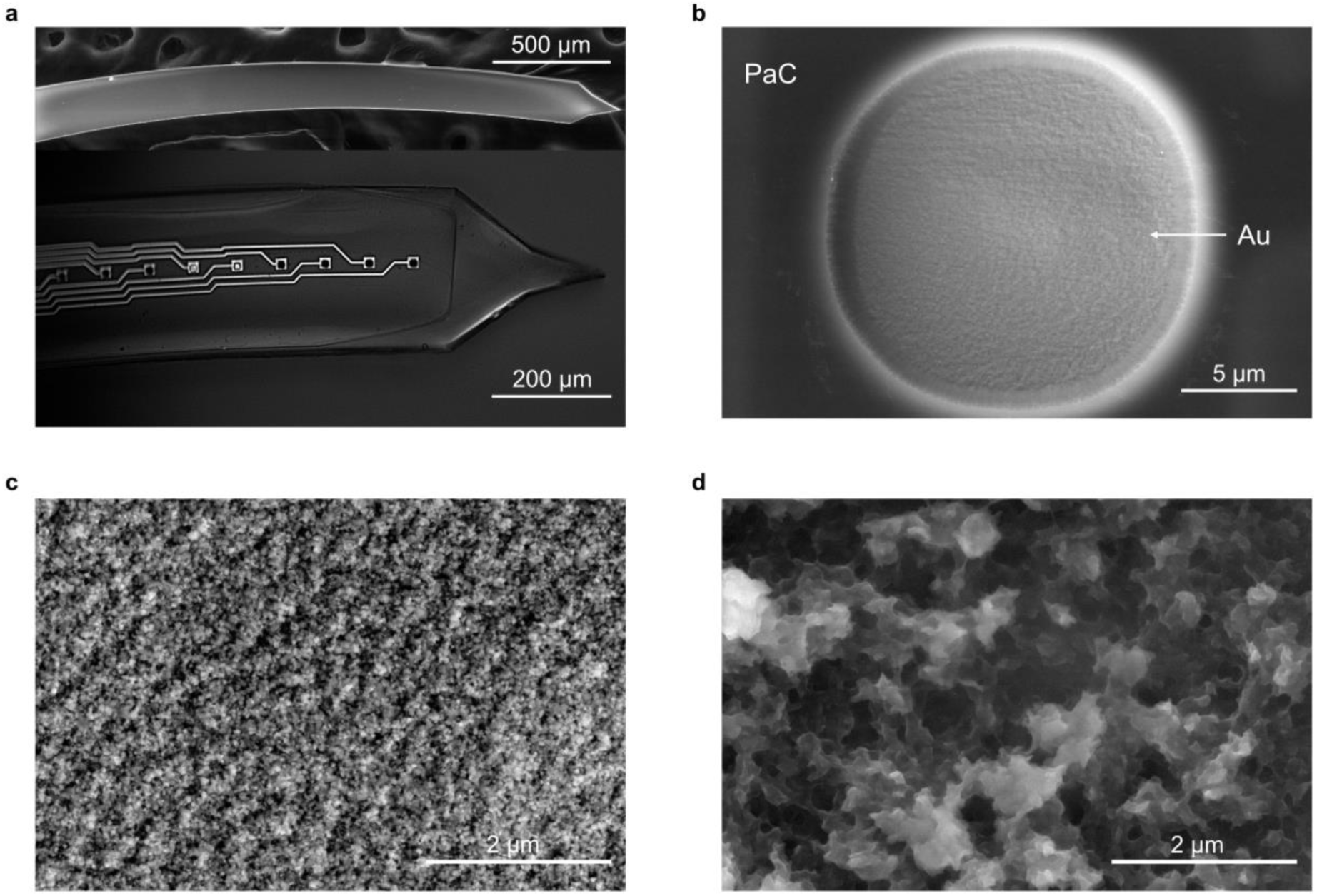
| Morphological characterization of the curved neural probe. **a** Optical micrograph of the 16-channel flexible microelectrode array integrated with the silk fibroin stiffener. (Top) Scanning electron microscope (SEM) image of the silk shuttle (backside view) showing the tapered, sharp tip geometry. **b** SEM image of an individual microelectrode site showing well-defined Parylene-C insulation boundaries. **c** SEM image of the Au electrode surface prior to PEDOT:BF_4_ deposition. **d** SEM image of the PEDOT:BF_4_-coated electrode surface, showing the characteristic nodular nanostructure. Scale bars are indicated on each panel.

### Optimization of PEDOT:BF_4_ Electrodeposition

To identify a PEDOT:BF_4_ coating suitable for chronic in vivo operation, we evaluated the electrochemical performance of electrodes deposited at different charge densities (50 to 1000 mC/cm^2^) (Fig. 3). Electrochemical impedance spectroscopy (EIS) (Fig. 3a) revealed a progressive decrease in impedance magnitude across the frequency range with increasing charge density, consistent with the formation of a thicker pseudocapacitive polymer film.

**Fig. 3.**
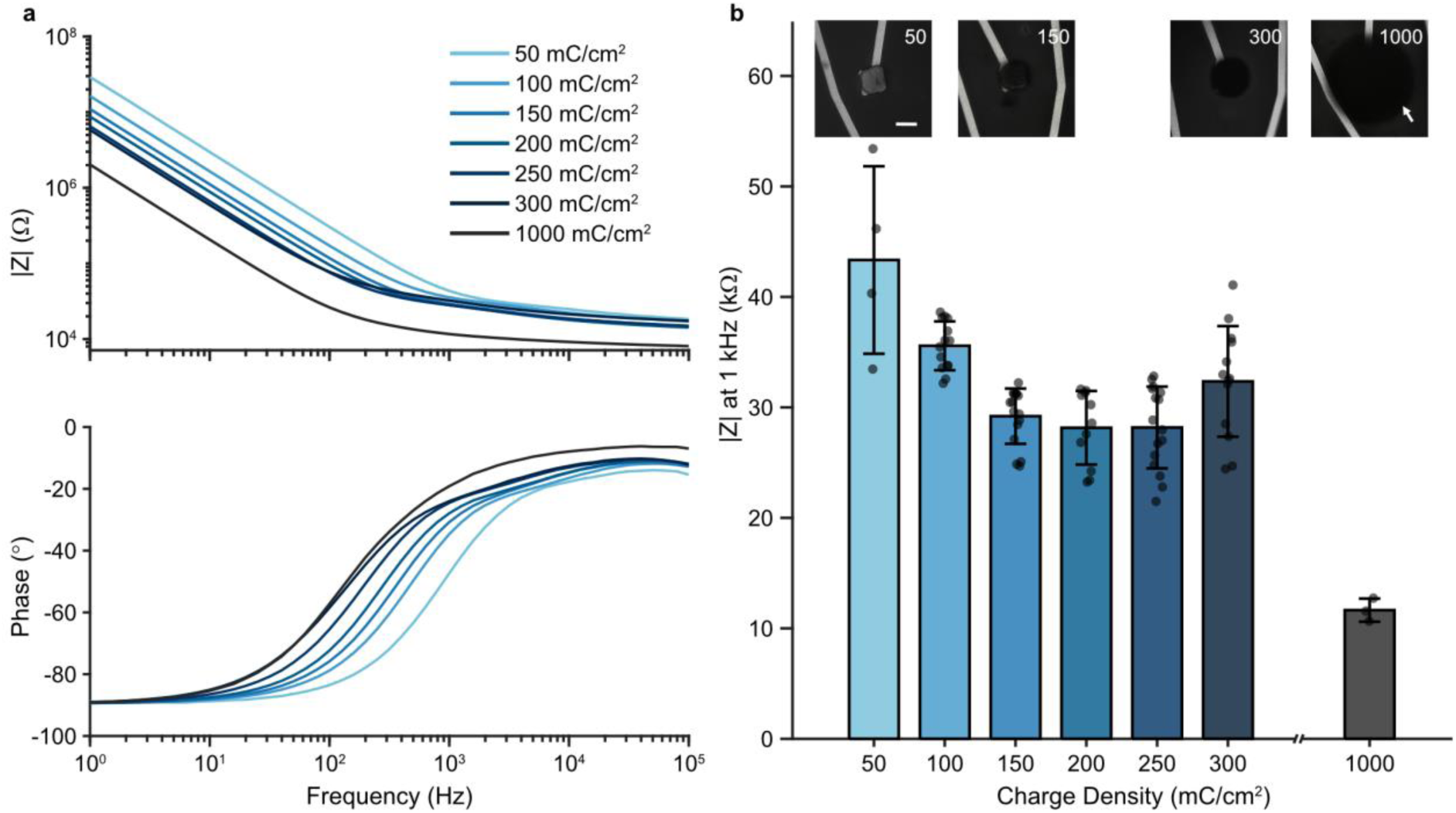
| Electrochemical and morphological optimization of PEDOT:BF_4_ deposition parameters. **a** Bode plots showing the impedance magnitude (|Z|, top) and phase angle (bottom) of microelectrodes following galvanostatic deposition at charge densities ranging from 50 to 1000 mC/cm^2^. Measurements were performed in 1x PBS. **b** Impedance magnitude at 1 kHz as at different deposition charge densites (bottom), with corresponding optical micrographs of the polymer coatings (top, scale bar = 10 µm). Increasing charge density led to extended deposition beyond the electrode area at high charge densities (white arrows). Bars represent mean values, error bars denote the standard deviation (s.d.), and individual points correspond to indiviual microelectrodes (*n* = 4 for 50 mC/cm², *n* = 3 for 1000 mC/cm², and *n* = 11–16 for all other groups).

The phase angle spectra further support this trend (Fig. 3a, bottom). Increasing the deposition charge density progressively shifted the −45° transition frequency from approximately 2 kHz to 150 Hz, extending the low-impedance resistive regime toward lower frequencies. This behaviour is consistent with the formation of a thicker pseudocapacitive PEDOT:BF_4_ coating.^29^

The impedance magnitude at 1 kHz was used as a benchmark to compare coatings (Fig. 3b). At low charge density (50 mC/cm^2^) electrodes exhibited high impedance (43.3 ± 8.5 kΩ), indicative of incomplete coverage. Increasing the deposition charge reduced impedance, reaching a plateau of 29.2 ± 2.5 kΩ at 150 mC/cm^2^ (*p* < 0.001). Beyond this point, further increases in charge density yielded minimal additional improvement,^30^ with no statistically significant difference between 150 mC/cm^2^ and 200 mC/cm^2^ (*p* = 0.99). Higher densities (up to 300 mC/cm^2^) resulted in increased variability without a corresponding improvement in average performance.

Morphological observations (Fig. 3b, top) support this electrochemical trend. At intermediate charge densities (150 mC/cm^2^) coatings were uniform and confined to the electrode area. In contrast, higher charge densities led to lateral overgrowth of the polymer beyond the electrode boundaries. Although very high charge densities (1000 mC/cm^2^) produced lower absolute impedance (11.6 ± 1.05 kΩ), this overgrowth compromises geometric definition and may compromise mechanical robustness during implantation.^19^ Based on this trade-off, a deposition charge density of 150 mC/cm^2^ was selected for all subsequent in vivo experiments, providing the best compromise between low impedance and controlled coating morphology.

### Electrochemical validation of PEDOT:BF_4_ coated electrodes and silk integration

Following optimization of the PEDOT:BF_4_ deposition parameters, the electrochemical performance of coated electrodes (150 mC/cm^2^) was compared with bare Au electrodes, and the effect of silk fibroin integration on electrode performance was evaluated (Fig. 4).

**Fig. 4.**
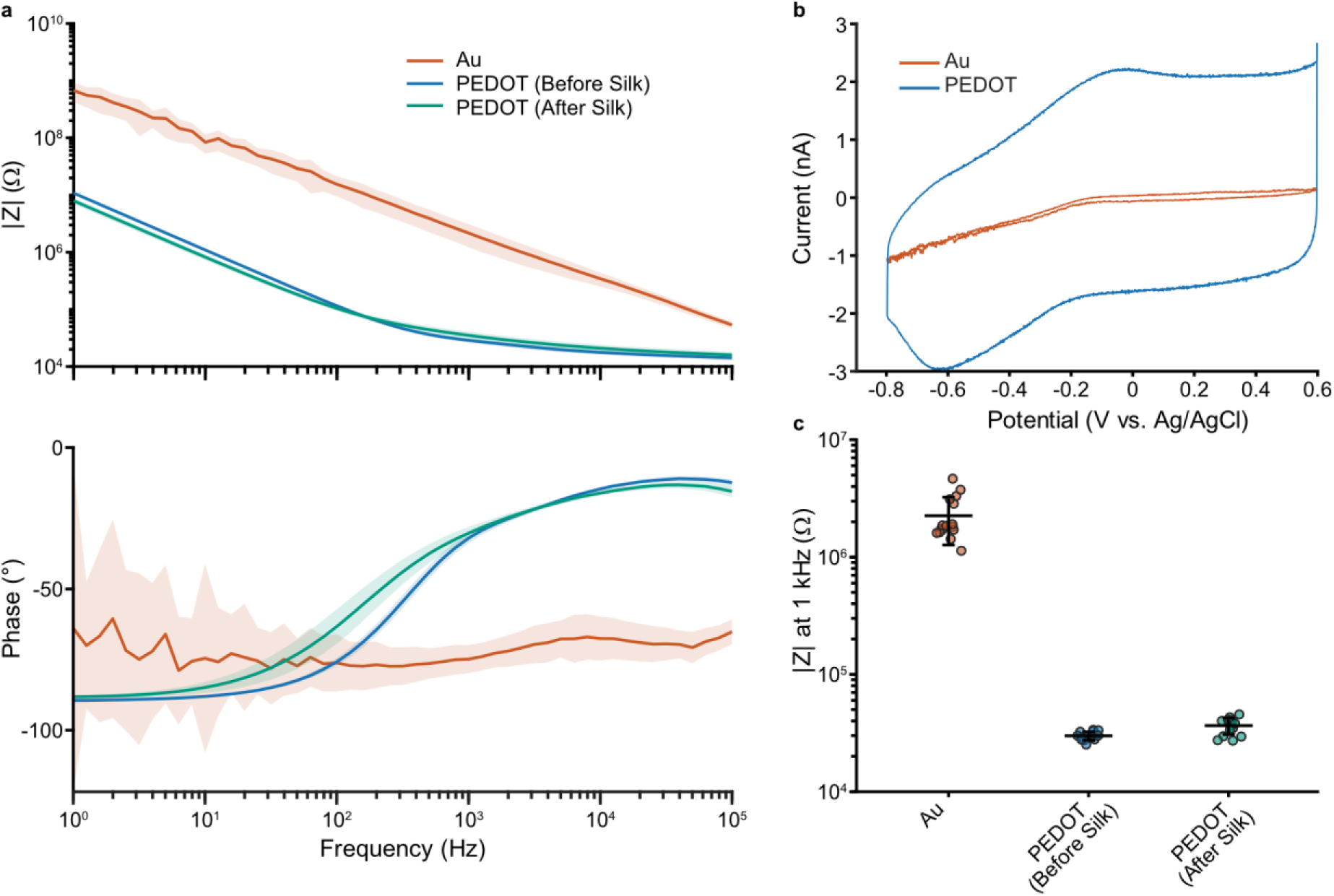
| Electrochemical performance of PEDOT:BF4-coated electrodes before and after integration with the silk stiffener. **a** Bode plots showing impedance magnitude (|Z|, top) and phase angle (bottom) for bare Au (*n* = 16), PEDOT:BF4 coated electrodes (*n* = 15), and PEDOT:BF4 after silk integration (*n* = 15). Shaded regions denote the standard deviation (s.d.). **b** Cyclic voltammetry (CV) of bare Au and optimized PEDOT:BF4 coated electrodes (scan rate 100 mV/s). **c** Impedance magnitude at 1 kHz for the three conditions plotted on a logarithmic scale. Individual points represent functional electrodes, while horizontal lines and error bars denote the standard deviation (s.d.). One-way ANOVA revealed no significant difference following silk integration (*p* > 0.05). Electrodeposition was performed at 150 mC/cm2. All measurements were performed in 1x PBS.

EIS analysis (Fig. 4a) revealed a substantial reduction in impedance across the full frequency range (1 Hz to 100 kHz) following PEDOT:BF_4_ deposition. At 1 kHz, the impedance decreased from 2.2 ± 0.9 MΩ for bare Au to 28.7 ± 2.3 kΩ for PEDOT-coated electrodes (*p* < 0.001) (Fig. 4c). The phase spectra further confirmed this improvement (Fig. 4a, bottom), evolving from the predominantly capacitive/blocking behaviour of bare Au toward a more pseudocapacitive interface.^31^ Cyclic voltammetry (Fig. 4b) further confirmed the enhanced electrochemical performance, with the cathodic charge storage capacity increasing approximately sixfold following PEDOT:BF₄ deposition.

Importantly, integration with the silk fibroin shuttle did not significantly affect electrode performance (Fig. 4c). The 1 kHz impedance remained statistically unchanged after silk integration (28.7 ± 2.3 kΩ to 35.1 ± 5.6 kΩ, *p* = 0.99), indicating that the silk-assisted implantation strategy preserves the electrochemical performance of the PEDOT:BF₄ interface.

### In vivo impedance evolution over time

To evaluate the evolution of the electrode–tissue interface over time, EIS analysis was performed repeatedly on the same implanted devices over a four-week period (Fig. 5). The impedance spectra (Fig. 5a) revealed an overall increase in magnitude following implantation, reflecting the dynamic evolution of the electrode-tissue interface.

**Fig. 5.**
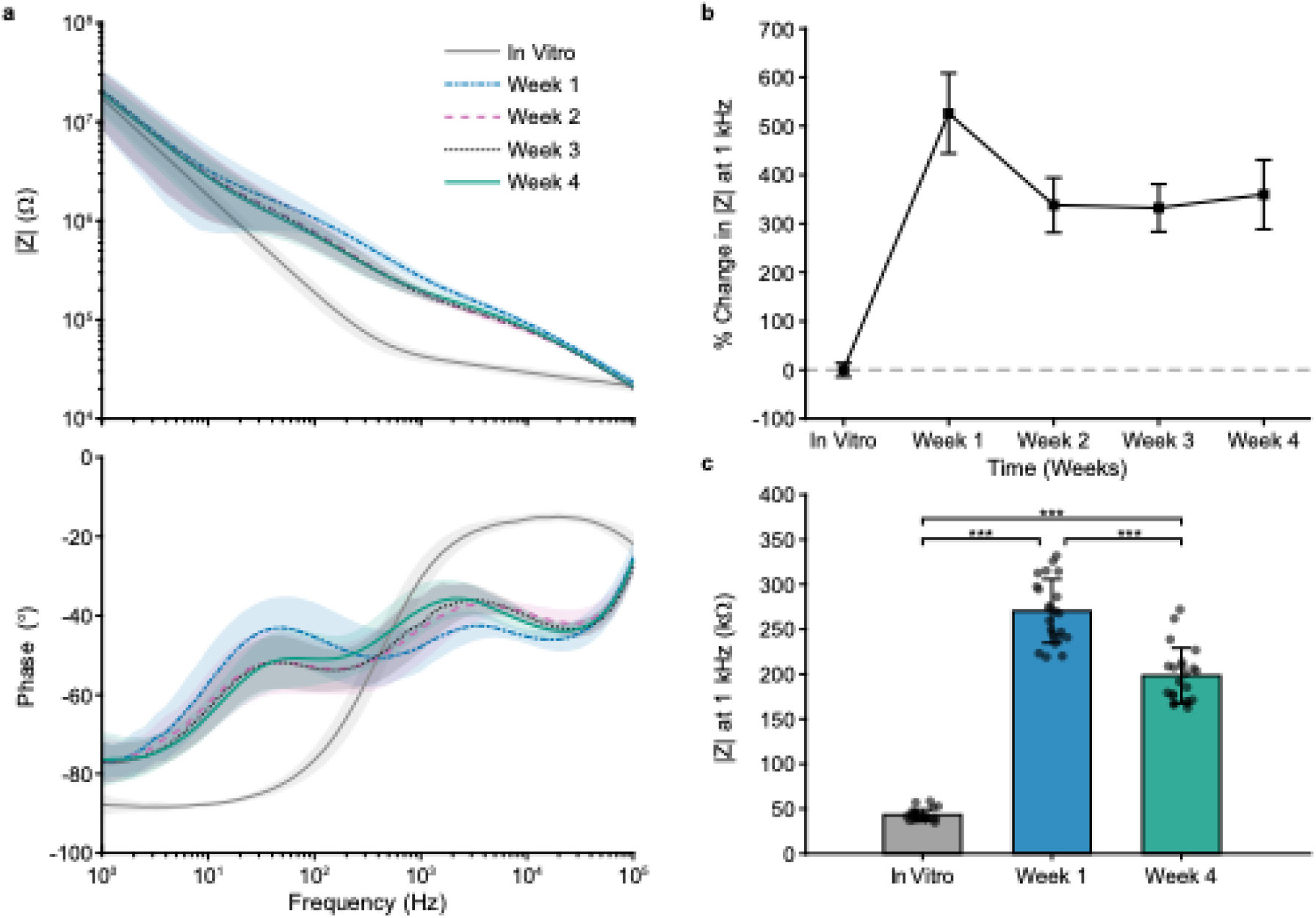
| In vivo evolution of the electrode-tissue interface over four weeks. **a** Bode plots showing the impedance magnitude (|Z|, top) and phase angle (bottom) from pre-implantation (in vitro) to Weeks 1–4 following implantation. Lines represent cohort mean values and shaded regions denote the standard deviation (s.d.) **b** Relative change in the impedance magnitude at 1 kHz over time. Squares indicate mean values and error bars denote s.d.; the dashed line indicates the pre-implantation baseline. **c** Absolute impedance magnitude at 1 kHz for representative time points (in vitro, Week 1, and Week 4). Bars and error bars denote the mean ±s.d. and individual points correspond to functional microelectrodes (*n* = 24 across *N* = 2 mice). Statistical significance was assessed using one-way ANOVA followed by Tukey’s post-hoc test, \*\*\**p* < 0.001.

To quantify these changes, the impedance at 1 kHz was monitored over time (Fig. 5b, c). Prior to implantation, PEDOT:BF_4_ coated microelectrodes exhibited a baseline impedance of 43.2 ± 6.1 kΩ. Following insertion into the ventral hippocampus, the impedance increased to 270.2 ± 35.5 kΩ at Week 1 (*p* < 0.001) (Fig. 5b, c), consistent with the acute tissue response.^32^ The impedance subsequently decreased to 189.0 ± 24.3 kΩ at Week 2 and stabilized through Week 4 (198.3 ± 30.7 kΩ), consistent with the stabilization of the chronic electrode–tissue interface.^33^ The decrease from the acute peak to Week 4 state is statistically significant (*p* < 0.001), while no significant variance was observed between Week 3 and Week 4 (*p* = 0.52), indicating stabilization of the recording interface.

The phase spectra (Fig. 5a) exhibited moderate changes in the mid-frequency region over time, consistent with the evolving tissue environment. Despite these variations, the low-frequency phase remained strongly capacitive (≈ −80° at 1 Hz) throughout the implantation period, indicating that the PEDOT:BF_4_ coating remained electrochemically active.

### Curved probe implantation and in vivo multisite recording in the vHP

Placement and recording capabilities of the curved microelectrode array were validated by targeting the CA1 sub-region of the ventral hippocampus (Fig. 6). Probe placement and trajectory were confirmed post-mortem using DAPI-stained coronal brain sections (Fig. 6a, b). The probes followed the predefined curved trajectories, enabling reproducible positioning of recording sites orthogonal to the pyramidal cell layer across CA1 and partially covering CA3. In one implantation (1 of 4), the probe trajectory terminated in ventral CA3 rather than CA1. Overall, these results demonstrate that curved microelectrode arrays can reproducibly access deep hippocampal regions while achieving an orientation that is difficult to obtain using conventional linear probes.

**Fig. 6.**
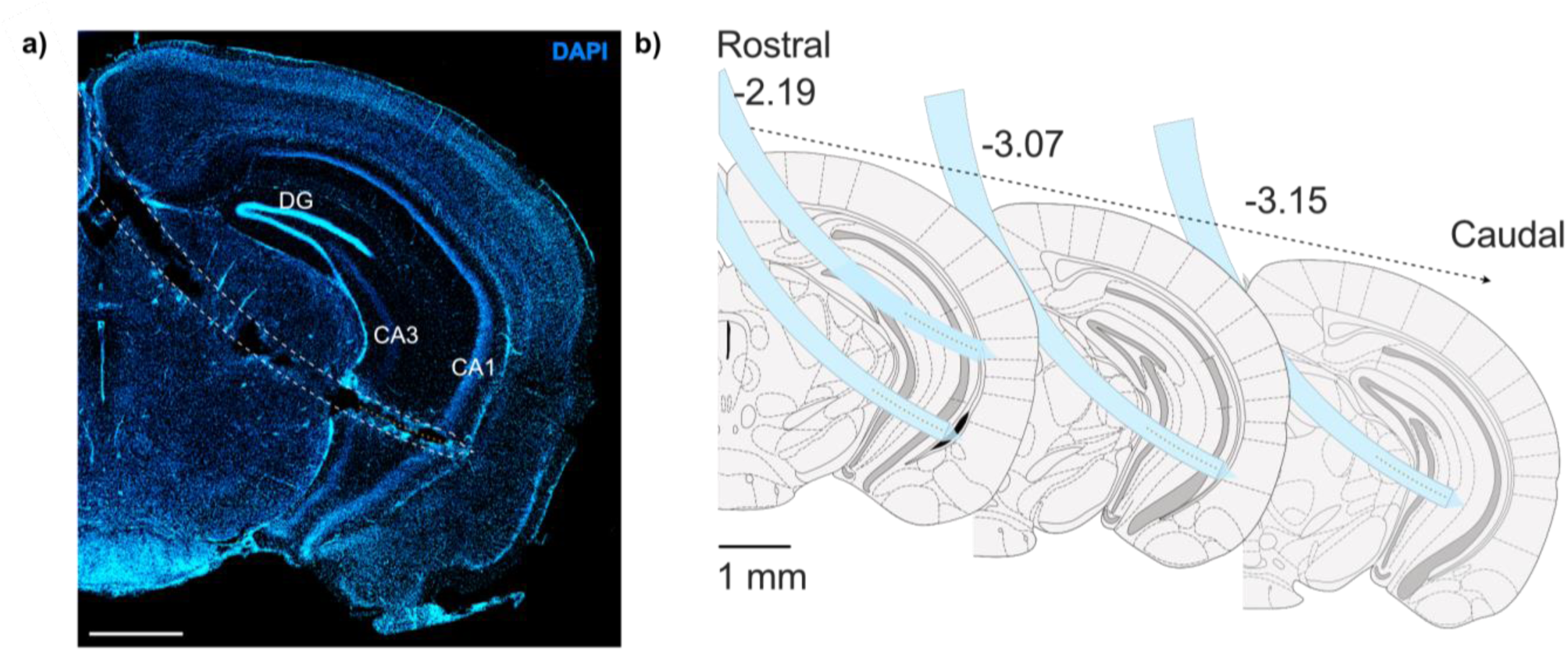
| Anatomical validation of curved probe placement in the ventral hippocampus. **a** Mouse brain section at the level of the ventral hippocampus, stained with the fluorescent cell nucleus marker DAPI, showing microelectrode track (delineated by the dotted lines). Hippocampal sub-regions CA1, CA3 and the dentate gyrus (DG) are indicated. **b** Schematics showing the position of the curved micro-electrode in four mice. Three different levels of the ventral hippocampus are shown along the rostro-caudal (front to back) axis of the mouse brain. Distances from the reference Bregma are indicated above each brain level (mm).

Acute in vivo recordings were performed to evaluate the functional performance of the curved neural probes. Ventral CA1 activity was recorded during rapid eye movement (REM) sleep, a behavioral state characterized by rhythmic 6-12 Hz LFP oscillations (Fig. 7a). Multisite recordings were subsequently used to compute current source density (CSD) profiles, which estimate the spatial distribution of transmembrane currents from extracellular voltage recordings and identify characteristic current sinks and sources associated with synaptic activity (Fig. 7b). The resulting patterns exhibited the expected sink–source alternations and enabled accurate identification of the relative positions of recording sites within hippocampal layers (Fig. 7b). These results demonstrate that the curved neural interface enables reliable layer-resolved recordings from the ventral hippocampus.

**Fig. 7.**
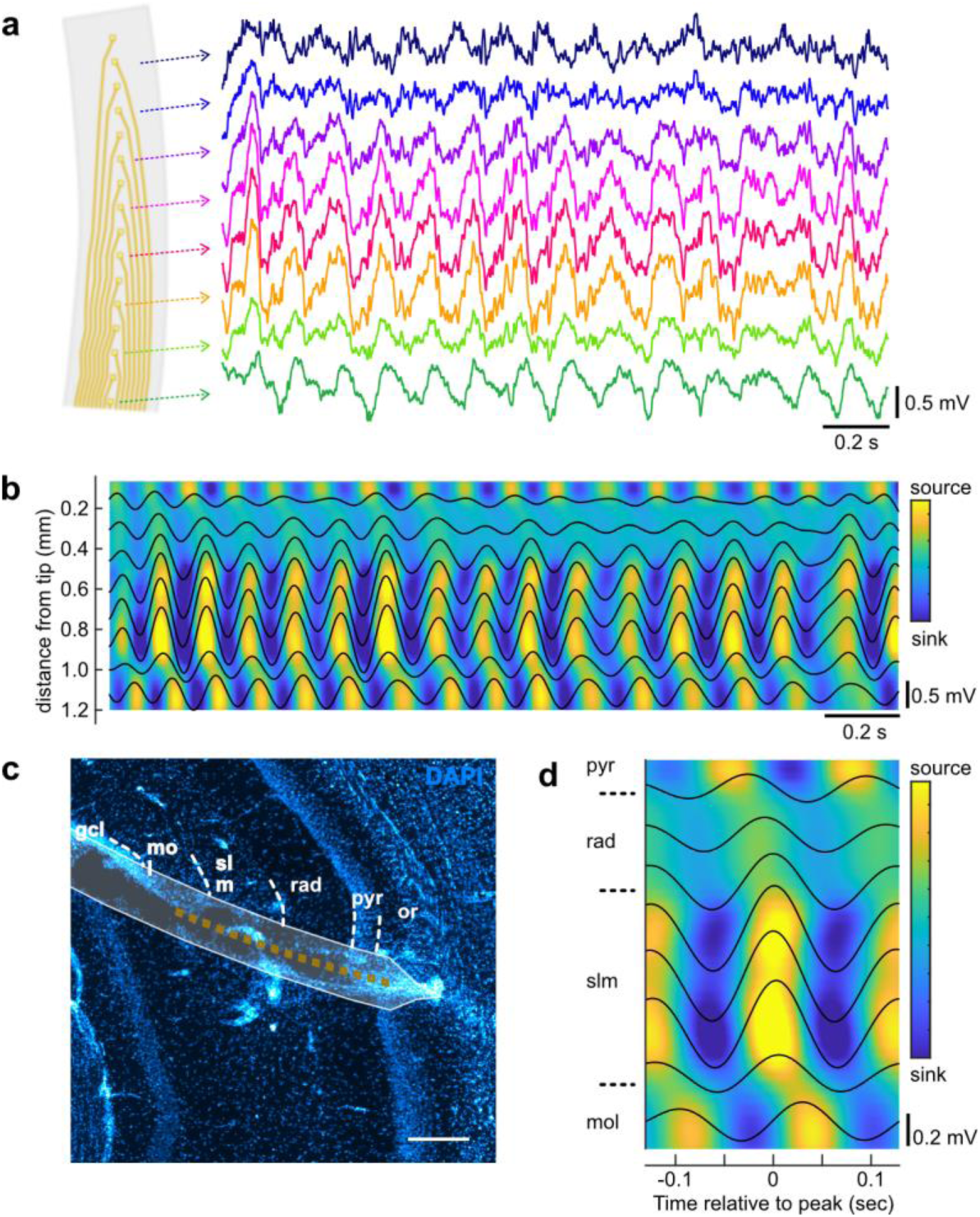
| Functional validation of curved neural interface in the ventral hippocampus. **a** Representative recordings from eight electrodes on the curved array acquired in the ventral hippocampus (vHP) during rapid-eye movement (REM) sleep. **b** Current source density (CSD) computed from recordings obtained in the CA1 region of the vHP during REM sleep. **c** Position of the curved array and estimated electrode location based on CSD analysis. Boundaries between ventral CA1 layers are indicated by the dotted lines. **d** CSD averaged to the peak of theta oscillations in stratum radiatum. Layer abbreviations: or: stratum oriens; pyr: stratum pyramidale; rad: stratum radiatum; sl m: stratum lacunosum moleculare; mo: molecular layer of the dentate gyrus; gcl: granule cell layer of the dentate gyrus. Scale bar: 250 µm.

## Discussion

Neural interfaces capable of resolving laminar activity in deep brain regions remain constrained by the geometric mismatch between linear probe architectures and the three-dimensional architectures of neural tissue.^2^ This limitation is particularly acute in brain regions with curved neuronal layers, such as the ventral hippocampus, where conventional linear probes struggle to achieve orthogonal alignment with the target anatomy. The curved microelectrode array and rotational insertion strategy presented here address this constraint by enabling anatomically matched access to deep brain structures.

A critical prerequisite for any neural probe is a high-quality electrode-tissue interface. Electrochemical characterization demonstrated that PEDOT:BF_4_ significantly improved performance, reducing impedance by two orders of magnitude at 1 kHz and providing a favorable balance between charge transfer and geometric confinement. While this optimization is essential, the central objective of this work was to translate these material properties into a functional in vivo neural interface.

Successful implantation in the ventral hippocampus demonstrates the feasibility of using pre-formed curved probes to target deep brain structures using a rotational insertion strategy. Histological analysis confirmed accurate probe placement within the target anatomy, while chronic recordings demonstrated stable acquisition of local field potentials over several weeks. These results show that anatomically matched probe geometry enables reliable layer-resolved current source density analysis in the ventral hippocampus.

Maintaining a stable electrode-tissue interface remains a major challenge in chronic neural recording.^34–36^ The impedance evolution observed in this study is consistent with the expected progression of the foreign body response following implantation. The initial increase in impedance at Week 1 likely reflects the acute response to insertion, including tissue disruption and early cellular activation at the interface.^37,38^ The transient presence of the silk stiffener may also contribute to these early interfacial changes.^39^ This initial increase was followed by a decrease and stabilization of impedance, rather than the progressive rise typically observed with rigid silicon probes. This behavior is consistent with the establishment of a stable chronic electrode-tissue interface, potentially facilitated by the mechanical compliance of the Parylene-C substrate, which may reduce motion-induced tissue damage.^38,40^ The absence of significant differences between Weeks 3 and 4 (*p* = 0.52) further supports stabilization of the interface over the time period investigated.

The phase spectra further support the stability of the system. While moderate changes in the mid-frequency regime indicate ongoing remodeling of the tissue environment,^41^ the consistently capacitive low-frequency response demonstrates that the PEDOT:BF_4_ coating remained electrochemically active throughout the implantation period.^6^

Although future studies comparing curved and linear probe architectures will be needed to isolate the contribution of geometry from substrate mechanics, the present results demonstrate that matching probe geometry to brain anatomy provides a practical strategy for layer-resolved recordings in complex three-dimensional neural structures.

## Conclusions

This work demonstrates that matching probe geometry to neural anatomy provides a practical strategy for accessing neuronal layers that are difficult to interrogate using conventional linear interfaces. By combining a predefined probe curvature with a transient silk fibroin stiffener, the proposed approach enables controlled alignment of the recording sites with curved neuronal layers while preserving mechanical compliance at the tissue interface.

The combination of a flexible Parylene-C substrate and a transient silk fibroin stiffener addresses the conflicting requirements of mechanical compliance after implantation and sufficient rigidity during insertion.

The resulting platform integrates low-impedance PEDOT:BF_4_-coated microelectrodes with stable in vivo performance, supporting laminar LFP recordings in deep hippocampal structures. The observed impedance stabilization and preserved electrochemical performance indicate that the interface remains functional under chronic conditions.

These results establish geometry-matched neural interfacing as a practical design strategy for flexible neural interfaces. Extending this approach to other anatomical targets and systematically comparing curved and linear probe designs will be important next steps toward defining the contribution of device geometry and expanding the design of neural interfaces for complex three-dimensional brain anatomy.

## Methods

### Design and fabrication of the curved microelectrode array

Curved probes were designed with a 7 mm radius of curvature to match the anatomical trajectory of the ventral hippocampus. Each device was fabricated on a Parylene-C (PaC) substrate and comprised sixteen gold (Au) microelectrodes (≈12 μm × 12 μm) separated by 75 μm (Fig. 1). Devices were fabricated on 4-inch glass wafers using standard photolithography and lift-off processes. A 2 μm layer of PaC film was deposited by chemical vapor deposition (SCS Labcoter 2, PDS 2010), followed by patterning of a Ti/Au (10/100 nm) layer for electrodes and interconnects. To pattern the Au, a bilayer photoresist was used, consisting of LOR5A (MicroChem, USA), spin-coated at 4000 RPM for 30 s and baked at 170 °C for 3 min, followed by OIR 674-11 (FUJIFILM, USA), spin-coated at 4000 RPM for 30 s, and baked at 90 °C for 90 s. The wafer was exposed to UV light (45 mJ/cm^2^) in a mask aligner (MA-6, Suss MicroTec, Germany) and developed for 75 s using AZ726 developer (Merck, Germany).

Prior to metal deposition, the wafer was treated with O_2_ reactive-ion etching (RIE) (100 W, 10 sccm, 1 min) to improve adhesion to Parylene-C. Au was deposited by electron beam evaporation electron (Thermionics, USA) and lift-off (Microposit Remover 1165, The Dow Chemical Company, USA). A second 2 μm PaC layer was added for insulation. The final probe geometry and electrode openings were defined using AZ9260 photoresist (AZ Electronic Materials, USA) spin-coated at 2400 RPM for 40 s, baked at 110 °C for 160 s, exposed to UV light (500 mJ/cm^2^) and developed in deionized water (DIW) and AZ400 K (AZ Electronic Materials, USA) at a 4:1 ratio for 3 min and 45 s. Patterning was achieved by dry etching with O_2_ (20 sccm) and CHF_3_ (2 sccm) at a pressure of 200 mTorr and a plasma power of 200 W for ∼25 min.^42^ The photoresist was removed by submerging the wafer in acetone for 1 min and rinsing with DIW.

Completed probes were released from the wafer by immersion in deionized water and transferred using fine-tipped brushes onto a transparent polyethylene terephthalate (PET) substrate for alignment and bonding. To improve the electrical connection of the 4 μm-thin parylene probe, a flat flexible cable (FFC, 0.25 mm pitch, 17 contacts, Molex) was electrically connected to the probe’s contact pads using an anisotropic conductive film (ACF) containing silver filler (128-32-F1, 1 mil thick, Creative Materials). A printed circuit board (PCB) was designed to interface with a flexible printed circuit (FPC) connector (0.25 mm pitch, 17 contacts, Molex) and a miniature connector (36-pin, NPD-36-DD-GS, Omnetics). The free end of the cable in the cable/probe assembly was inserted into the FPC connector, allowing this setup to connect with external circuits for conducting electrochemical and neural recording experiments.

### Electrochemical cleaning and electrodeposition of PEDOT:BF_4_

Electrochemical experiments were performed using a potentiostat (BioLogic) and a three-electrode glass cell housed in a Faraday cage (BioLogic). Measurements were carried out in deaerated solutions under a continuous nitrogen flow. The microelectrodes of the Parylene-C probe served as the working electrode, while a platinum coil and an Ag/AgCl (3 M NaCl) (RE-5B) electrode were used as the counter and reference electrodes, respectively.

All Au microelectrodes (≈ 144 μm^2^) were electrochemically cleaned and restructured by cyclic voltammetry in 0.2 M H_2_SO_4_ by sweeping the potential from 0 V to +1.6 V vs Ag/AgCl at 0.3 V/s up to 50 cycles, followed by EIS characterization. PEDOT films were subsequently galvanostatically electrodeposited from an acetonitrile solution containing 0.02 M EDOT and 0.12 M TEABF_4_. Electropolymerization was performed at a current density of 1 mA/cm^2^ for deposition times ranging from 50 s to 16 min, yielding polymerization charge densities ranging from 50 to 1000 mC/cm^2^.

Electrode morphology was characterized by scanning electron microscopy (SEM) using a Quattro variable pressure SEM (Thermo Fisher) with accelerating voltages ranging from 10 kV to 20 kV, a current of 100 pA or 140 pA, and a pressure of 80 Pa. Energy dispersive X-ray spectroscopy (EDS) was used to confirm the cleanliness of the Au surface.

### In vitro electrochemical characterization

In vitro electrochemical characterization studies were conducted using the same setup described for electrodeposition. Cyclic voltammetry (CV) and electrochemical impedance spectroscopy (EIS) were performed at room temperature in phosphate buffer saline (PBS, Sigma Aldrich, 0.01 M phosphate buffer, 0.0027 M potassium chloride, and 0.137 M sodium chloride, pH 7.4, at 25 °C). EIS measurements were performed at open circuit potential using a 10 mV (r.m.s.) sinusoidal perturbation over a frequency range of 1 Hz to 100 kHz.

For the CV measurements, the electrode potential was swept between -0.8 V and +0.6 V (vs. Ag/AgCl) for 5 cycles at a scan rate of 0.1 V/s. The cathodic charge storage capacity (CSCc) was calculated by integrating the cathodic current over the full potential range using the trapezoidal rule, normalizing by the scan rate and the geometric surface area of the electrode. To minimize the influence of inter-cycle variations, CSCc values were calculated from the average response of the final four cycles.

### Silk stiffener preparation and probe assembly

Transient silk fibroin stiffeners were fabricated to provide temporary mechanical support during implantation. Aqueous silk fibroin solution was regenerated from *Bombyx Mori* cocoons following established protocols ^43–45^ and concentrated to 10% w/v before storage at 4 °C. A PDMS mold was fabricated from a patterned silicon master using deep reactive-ion etching (DRIE) to define the curved geometry. The mold was designed to produce stiffeners with a sharp tip, to facilitate tissue penetration, and a cross-sectional area of approximately 300 × 300 μm.

For device assembly, the Parylene-C probe was positioned in the PDMS mold with the electrode side facing the mold to keep the recording sites exposed. The silk fibroin solution was then cast onto the probe and allowed to dry at ambient temperature. After drying, the silk-integrated probe was removed from the mold using forceps and lightly clamped between silicone foam for at least 12 h to preserve its geometry. The assembled devices were subsequently stored under ambient conditions for 12–24 h prior to implantation.

### Animal models and surgical procedures

All animal procedures were conducted in accordance with Canadian Council of Animal Care guidelines and approved by the CHU Sainte-Justine Research Center Animal Ethics Board (CIBPAR). Four C57BL/6J mice (Jackson Laboratory stock no. 000664, 2–4 months old) were used in this study and implanted in the ventral hippocampus. Animals were maintained under controlled environmental conditions (23 °C, 12 h light/dark cycle), with food and water provided ad libitum.

On the day of the surgery, anesthesia was induced with 5% isoflurane in O_2_ and maintained at 2% during the procedure. Animals were positioned in a stereotaxic frame, and body temperature was maintained at 37.5 °C using a heating pad.

A rotating implantation system was developed to insert the curved probes along their predefined trajectory. The system consisted of a custom probe holder attached to a rotational manipulator mounted on a standard stereotaxic arm, positioning the probe centerline 7 mm from the axis of rotation. Target coordinates were determined from a reference brain atlas, and the probe’s geometry was overlaid using AutoCAD to identify the optimal entry point. Following identification of the Bregma, a craniotomy was performed at the entry coordinates of the ventral hippocampus (−3.08 mm anterior/posterior, 0.39 mm medial/lateral). A stainless-steel screw placed above the cerebellum served as a ground/reference electrode, and a second posterior screw was used for mechanical anchoring.

The probe tip was positioned at the cortical surface, and the dura was opened prior to implantation. The probe was then rotated by 63°, reaching its final target coordinates: −3.08 mm anterior/posterior, 3.80 mm medial/lateral, and −3.79 mm dorso/ventral. The external part of the electrodes was secured to the skull using dental cement (Paterson Dental) and the flexible cable was attached to the PCB through the FPC connector, before final encapsulation with additional dental cement.

### In vivo electrochemical impedance spectroscopy (EIS)

To evaluate the longitudinal electrochemical stability of the implanted arrays, in vivo EIS was performed weekly over a 4-week period. During measurements, mice were placed in a non-conductive plastic enclosure shielded by a custom copper-mesh Faraday cage.

Electrical connection to the potentiostat was established by mating a 36-pin female dual-row wire connector (A79029-001, Omnetics) to the skull-mounted assembly. Measurements were conducted in a two-electrode configuration: the implanted microelectrodes served as the working electrodes (WE), while the cerebellar stainless-steel screw functioned as the combined counter and reference electrode (CE/RE). The potentiostat, Faraday cage, and earth ground were connected to a single common grounding point to prevent earth loops. Impedance spectra were acquired using a PalmSens4 potentiostat (PalmSens BV) communicating via Bluetooth. EIS was conducted at OCP, recorded for 60 s, followed by a 5 s equilibration period. An AC sinusoidal perturbation of 10 mV (r.m.s.) was applied, and the frequency was swept logarithmically from 100 kHz to 1 Hz (10 points per decade). Current measurement limits were set between 1 nA and 10 μA.

### In vivo acute recordings and histology

Neural signals were acquired using a 32-channel Open Ephys acquisition system,^47^ at 30 kSamples/s and bandpass filtered online between 0.1 Hz - 14 kHz. Recordings were performed in freely behaving animals during the normal sleep-wake cycles (awake, rapid eye movement [REM] sleep and non-REM sleep). Neural data were subsequently processed offline using custom Matlab scripts. Signals were filtered between 3 and 12 Hz to isolate theta oscillations, and periods of REM sleep were identified visually. Current source density (CSD) profiles were computed using the iCSD method (Petterson et al. J Neuroscience Methods 2006). For theta-locked average CSD analysis, the LFP recorded in the stratum radiatum was used as reference signal. Theta oscillation peaks were detected and theta peak locked averages for each channel were computed prior to CSD computation.

At the end of the experiments, animals were deeply anesthetized with a ketamine/xylazine/acepromazine cocktail (80/12.5/2.5 mg/kg) via intraperitoneal injection and transcardially perfused with phosphate-buffered saline (PBS). Brains were extracted and fixed in 4% w/v paraformaldehyde for one week to preserve the electrode tracks. The samples were sectioned into 25 μm slices using a Vibratome (Leica VT 1200S) and processed following standard histological technique. Processed sections were mounted with 4,6-diamidino-2-phenylindole (DAPI) Fluoromount-G (SouthernBiotech) and stored in the dark at 4 °C prior to imaging. Fluorescence microscopy was used to confirm probe trajectories and electrode locations.

### Data processing and statistical analysis

Electrochemical data processing, visualization and statistical data analysis were performed using custom MATLAB scripts. For the in vitro optimization of PEDOT:BF_4_ deposition conditions, statistics were calculated by pooling all functional microelectrodes. Sample sizes ranged from *n* = 11–16 functional replicates for deposition charge densities between 100 and 300 mC/cm², with *n* = 4 and *n* = 3 for 50 mC/cm² and 1000 mC/cm², respectively.

For longitudinal in vivo analysis, impedance magnitude (|Z|) and phase angle were extracted at each frequency and calculated by pooling all functional microelectrodes (*n* = 24) across the biological replicates (mice, *N* = 2) for each time point. Relative impedance changes were calculated by normalizing the 1 kHz impedance of implanted electrodes to their corresponding pre-implantation PEDOT:BF_4_ values.

Variance is visualized using bar charts and longitudinal trend lines overlaid with the individual data points of the functional replicates. Statistical significance of the interface evolution across all five temporal stages (in vitro and Weeks 1 through 4), a one-way analysis of variance (ANOVA) was performed on |Z| and the phase angle at 1 kHz. Because this longitudinal study required the identification of specific temporal transitions— such as the acute inflammatory peak versus the chronic plateau—Tukey’s Honestly Significant Difference (HSD) post-hoc test was applied following the ANOVA. This approach strictly controls the family-wise error rate across all pairwise multiple comparisons. All data are reported as the mean ± standard deviation (s.d.), and a *p*-value of *p* < 0.05 was considered statistically significant.

## Data availability

The datasets generated and analysed during the current study are publicly available in the Zenodo repository at https://doi.org/10.5281/zenodo.21373465.

## Acknowledgements

We thank Dr. Elke Küster-Schöck and Dr. Vanesa Jimenez Amilburu from the Centre Hospitalier Universitaire Sainte-Justine Imaging Center for their assistance with microscopy, and Véronique Pelletier and the animal facility team for their support with animal colony maintenance. This work was supported by the New Frontiers in Research Fund (NFRF) Exploration program (NFRFE-2022-00447) awarded to B.A. and F.C., and by a Natural Sciences and Engineering Research Council of Canada (NSERC) Discovery Grant (RGPIN-2025-05607) awarded to F.C. Equipment and infrastructure used in this study were funded by the Canada Foundation for Innovation (CFI). J.H. acknowledges financial support from the TransMedTech Institute through a doctoral scholarship funded by the Canada First Research Excellence Fund (CFREF).

## Author contributions

B.A. and F.C. conceived the study, supervised the research, and secured funding. J.H. designed and fabricated the neural probes, performed the PEDOT:BF₄ electrodeposition, electrochemical and morphological characterization, analyzed the data, performed the statistical analysis, prepared the figures, and drafted the manuscript. G.D. performed the animal surgeries, in vivo electrophysiological recordings, and contributed to histology, imaging, data analysis, and figure preparation. L.P. contributed to the design of the in vivo experiments and assisted with the animal surgeries, electrophysiological recordings, histology, imaging, and data analysis. Y.M.-P. contributed to the electrochemical characterization and data analysis. M.W. contributed to the microfabrication of the neural probes. J.H., B.A., and F.C. wrote the manuscript with input from all authors. All authors discussed the results, revised the manuscript, and approved the final version.

## References

1. Renshaw, B., Forbes, A. & Morison, B. R. Activity of isocortex and hippocampus: electrical studies with micro-electrodes. J. Neurophysiol. 3, 74–105 (1940).

2. Royer, S., Sirota, A., Patel, J. & Buzsáki, G. Distinct representations and theta dynamics in dorsal and ventral hippocampus. J. Neurosci. 30, 1777–1787 (2010).

3. Buzsáki, G. Theta oscillations in the hippocampus. Neuron 33, 325–340 (2002).

4. Ekstrom, A. D. et al. Human hippocampal theta activity during virtual navigation. Hippocampus 15, 881–889 (2005).

5. Strange, B. A., Witter, M. P., Lein, E. S. & Moser, E. I. Functional organization of the hippocampal longitudinal axis. Nat. Rev. Neurosci. 15, 655–669 (2014).

6. Rivnay, J., Wang, H., Fenno, L., Deisseroth, K. & Malliaras, G. G. Next-generation probes, particles, and proteins for neural interfacing. Sci. Adv. 3, e1601649 (2017).

7. Hong, G. & Lieber, C. M. Novel electrode technologies for neural recordings. Nat. Rev. Neurosci. 20, 330–345 (2019).

8. Ferro, M. D. et al. NeuroRoots, a bio-inspired, seamless brain machine interface for long-term recording in delicate brain regions. AIP Adv. 14, (2024).

9. Zhao, Z. et al. Ultraflexible electrode arrays for months-long high-density electrophysiological mapping of thousands of neurons in rodents. *Nat*. Biomed. Eng. 7, 520–532 (2023).

10. Lacour, S. P., Courtine, G. & Guck, J. Materials and technologies for soft implantable neuroprostheses. Nat. Rev. Mater. 1, 1–14 (2016).

11. Du, Z. J. et al. Ultrasoft microwire neural electrodes improve chronic tissue integration. Acta Biomater. 53, 46–58 (2017).

12. Cui, X. & Martin, D. C. Electrochemical deposition and characterization of poly (3, 4-ethylenedioxythiophene) on neural microelectrode arrays. Sensors Actuators B Chem. 89, 92–102 (2003).

13. Dimov, I. B., Moser, M., Malliaras, G. G. & McCulloch, I. Semiconducting polymers for neural applications. Chem. Rev. 122, 4356–4396 (2022).

14. Bodart, C. et al. Electropolymerized poly (3, 4-ethylenedioxythiophene)(PEDOT) coatings for implantable deep-brain-stimulating microelectrodes. ACS Appl. Mater. Interfaces 11, 17226–17233 (2019).

15. Hagler, J. et al. Electrodeposited PEDOT: BF4 coatings improve impedance of chronic neural stimulating probes in vivo. Adv. Mater. Interfaces 9, 2201066 (2022).

16. Castagnola, V. et al. Parylene-based flexible neural probes with PEDOT coated surface for brain stimulation and recording. Biosens. Bioelectron. 67, 450–457 (2015).

17. Fanselow, M. S. & Dong, H.-W. Are the dorsal and ventral hippocampus functionally distinct structures? Neuron 65, 7–19 (2010).

18. Boehler, C., Aqrawe, Z. & Asplund, M. Applications of PEDOT in Bioelectronic Medicine. Bioelectron. Med. 2, 89–99 (2019).

19. Ludwig, K. A. et al. Poly (3, 4-ethylenedioxythiophene)(PEDOT) polymer coatings facilitate smaller neural recording electrodes. J. Neural Eng. 8, 014001 (2011).

20. Venkatraman, S. et al. In Vitro and In Vivo Evaluation of PEDOT Microelectrodes for Neural Stimulation and Recording. IEEE Trans. Neural Syst. Rehabil. Eng. 19, 307–316 (2011).

21. Hassler, C., Boretius, T. & Stieglitz, T. Polymers for neural implants. J. Polym. Sci. Part B Polym. Phys. 49, 18–33 (2011).

22. Gao, W. et al. Revolutionizing brain‒computer interfaces: overcoming biocompatibility challenges in implantable neural interfaces. J. Nanobiotechnology 23, 498 (2025).

23. Buzsáki, G. Large-scale recording of neuronal ensembles. Nat. Neurosci. 7, 446–451 (2004).

24. Nicholson, C. & Freeman, J. A. Theory of current source-density analysis and determination of conductivity tensor for anuran cerebellum. J. Neurophysiol. 38, 356–368 (1975).

25. Barnes, M. C., Kim, D.-Y., Ahn, H. S., Lee, C. O. & Hwang, N. M. Deposition mechanism of gold by thermal evaporation: approach by charged cluster model. J. Cryst. Growth 213, 83– 92 (2000).

26. Boehler, C., Vieira, D. M., Egert, U. & Asplund, M. NanoPt—a nanostructured electrode coating for neural recording and microstimulation. ACS Appl. Mater. Interfaces 12, 14855– 14865 (2020).

27. Yang, J. & Martin, D. C. Impedance spectroscopy and nanoindentation of conducting poly (3, 4-ethylenedioxythiophene) coatings on microfabricated neural prosthetic devices. J. Mater. Res. 21, 1124–1132 (2006).

28. Castagnola, V., Bayon, C., Descamps, E. & Bergaud, C. Morphology and conductivity of PEDOT layers produced by different electrochemical routes. Synth. Met. 189, 7–16 (2014).

29. Boehler, C., Carli, S., Fadiga, L., Stieglitz, T. & Asplund, M. Tutorial: guidelines for standardized performance tests for electrodes intended for neural interfaces and bioelectronics. Nat. Protoc. 15, 3557–3578 (2020).

30. Baek, S., Green, R. A. & Poole-Warren, L. A. The biological and electrical trade-offs related to the thickness of conducting polymers for neural applications. Acta Biomater. 10, 3048– 3058 (2014).

31. Lecomte, A., Degache, A., Descamps, E., Dahan, L. & Bergaud, C. In vitro and in vivo biostability assessment of chronically-implanted Parylene C neural sensors. Sensors Actuators B Chem. 251, 1001–1008 (2017).

32. Prasad, A. & Sanchez, J. C. Quantifying long-term microelectrode array functionality using chronic in vivo impedance testing. J. Neural Eng. 9, 26028 (2012).

33. Williams, J. C., Hippensteel, J. A., Dilgen, J., Shain, W. & Kipke, D. R. Complex impedance spectroscopy for monitoring tissue responses to inserted neural implants. J. Neural Eng. 4, 410–423 (2007).

34. Siwakoti, U., Jones, S. A., Kumbhare, D., Cui, X. T. & Castagnola, E. Recent Progress in Flexible Microelectrode Arrays for Combined Electrophysiological and Electrochemical Sensing. Biosensors vol. 15 100 at 10.3390/bios15020100 (2025).

35. Williams, N. P. et al. In vivo microelectrode arrays for neuroscience. Nat. Rev. Methods Prim. 5, 31 (2025).

36. Lv, S. et al. Long-term stability strategies of deep brain flexible neural interface. *npj Flex*. Electron. 9, 40 (2025).

37. Salatino, J. W., Ludwig, K. A., Kozai, T. D. Y. & Purcell, E. K. Glial responses to implanted electrodes in the brain. *Nat*. Biomed. Eng. 1, 862–877 (2017).

38. Polikov, V. S., Tresco, P. A. & Reichert, W. M. Response of brain tissue to chronically implanted neural electrodes. J. Neurosci. Methods 148, 1–18 (2005).

39. Sommakia, S., Rickus, J. L. & Otto, K. J. Effects of adsorbed proteins, an antifouling agent and long-duration DC voltage pulses on the impedance of silicon-based neural microelectrodes. in 2009 Annual International Conference of the IEEE Engineering in Medicine and Biology Society 7139–7142 (2009). doi:10.1109/IEMBS.2009.5332456.

40. Sillay, K. A. et al. Long-Term Surface Electrode Impedance Recordings Associated with Gliosis for a Closed-Loop Neurostimulation Device. Ann. Neurosci. 25, 289–298 (2019).

41. Hazelgrove, B. et al. Electrochemical impedance spectroscopy in vivo for neurotechnology and bioelectronics. Nat. Rev. Electr. Eng. 2, 110–124 (2025).

42. Pas, J. et al. A bilayered PVA/PLGA-bioresorbable shuttle to improve the implantation of flexible neural probes. J. Neural Eng. 15, 065001 (2018).

43. Romero, I. S., Schurr, M. L., Lally, J. V, Kotlik, M. Z. & Murphy, A. R. Enhancing the interface in silk–polypyrrole composites through chemical modification of silk fibroin. ACS Appl. Mater. Interfaces 5, 553–564 (2013).

44. Murphy, A. R., John, P. S. & Kaplan, D. L. Modification of silk fibroin using diazonium coupling chemistry and the effects on hMSC proliferation and differentiation. Biomaterials 29, 2829–2838 (2008).

45. Hagler, J. R., Peterson, B. N., Murphy, A. R. & Leger, J. M. Performance of silk-polypyrrole bilayer actuators under biologically relevant conditions. J. Appl. Polym. Sci. 136, 46922 (2019).

46. Paxinos, G. & Franklin, K. B. J. Paxinos and Franklin’s the Mouse Brain in Stereotaxic Coordinates. (Academic press, 2019).

47. Siegle, J. H. et al. Open Ephys: an open-source, plugin-based platform for multichannel electrophysiology. J. Neural Eng. 14, 45003 (2017).

